# An interpretable peptide–HLA model emergently learns binding energetics and structure

**DOI:** 10.64898/2026.09.08.750216

**Authors:** Sully F. Chen, Robert J. Steele, Eric K. Oermann

## Abstract

The range of peptides a human leukocyte antigen (HLA) binds and displays modulates immune response and therefore underpins vaccine design, neoantigen discovery, autoimmunity, transplantation, and hypersensitivity reactions. Modern predictors of peptide–HLA (pMHC) binding and presentation are remarkably accurate, but they are black boxes; their internal computations are opaque and post-hoc explanatory methods lack guarantees of attribution. Here we introduce *LAtent Motif INteraction Aggregation* (LAMINA), an architecture whose prediction is, by construction, interpretable and attributable. LAMINA embeds every gapless subsequence of the HLA pseudosequence and of the candidate peptide as a learned “soft” motif, scores every HLA–peptide-motif pair, and lastly aggregates those scores to produce a prediction. Despite having only 4.7 million parameters and training in about ten hours on a single desktop workstation, LAMINA matches or exceeds state-of-the-art predictors on held-out binding-affinity regression and is competitive on rank correlation. Strikingly, the model states correlate with interaction energies and other structural metrics in structurally characterized pMHC complexes, despite training exclusively on sequence data alone.

## 1 Introduction

Class I major histocompatibility complex (MHC) molecules continuously sample the intracellular proteome and display short peptides on the cell surface for inspection by CD8^+^ T-cells^1^. The structural basis of this display was established by the first crystal structures of HLA-A2, which revealed a closed groove formed by two *α*-helices over an eight-stranded *β*-sheet, with pockets that clamp the peptide termini^2,3^. Pool sequencing of naturally eluted peptides later showed that each allele imposes an allele-specific sequence motif, in which a small number of “anchor” positions, classically P2 and the C-terminus, are strongly constrained while the intervening positions are largely free to vary ^4–6^. This combination of a rigid groove and a permissive middle is what makes the problem both tractable and hard: a handful of positions carry most of the specificity, but the remaining positions, peptide length, and the tens of thousands of known class I alleles^7^ interact in ways that no single motif captures.

The stakes for accurate interaction prediction are high; peptide–HLA (pMHC) prediction is the first computational filter in essentially every neoantigen-discovery pipeline^8,9^, and shapes which epitopes are chosen for personalized cancer vaccines that are now undergoing clinical trials^10–13^. These same predictions inform infectious-disease vaccine design, the interpretation of HLA associations in autoimmunity, and the mechanistic understanding of HLA-modulated drug hypersensitivity ^14,15^. In each of these settings, immunologists desire to know both the predicted interaction strength and *which residues* drive the score, so that candidates can be modified, mechanisms proposed, or unexpected predictions further investigated.

Computational pMHC prediction has a remarkably long history. Early methods used explicit motif logic: side-chain coefficient tables ranked HLA-A2 binders by modeling positions as independent^16^, curated motif databases codified allele-specific anchors^17^, and position-specific scoring matrices with stabilizing regularization^18,19^. Neural networks eventually replaced the independence assumption with learned higher-order structure, first for single alleles^20,21^ and then pan-allelic so that a single model could generalize across alleles and even across species^22,23^. Critically, the arrival of large-scale mass-spectrometry immunopeptidomics data^24–27^ produced the current generation of predictors, which jointly model *in vitro* binding affinity and naturally eluted ligands and deconvolve multi-allelic samples during training^28–35^. More recent work has incorporated deep learning techniques spanning graph neural networks, transformer architectures, and, most recently, protein language models ^36–42^, combined with in-silico structure-based modeling^43–45^. Benchmarks consistently place the best of these tools at a high level of accuracy ^46,47^.

These modern pMHC predictors then are often quite complex: multi-layer networks with nonlinearities, attention/graphs, or other complex operations making their inner workings challenging to interpret. Post-hoc explanations such as surrogate local models^48^, Shapley-value approximations ^49^, path-integrated gradients ^50^, reference-based activation differences ^51^, or simply inspecting attention maps all offer limited visibility into model dynamics. These tools are sometimes insightful, but often only weakly related to the underlying computation. For high-stakes biological decisions or expensive in vivo research pipelines, a model would ideally be interpretable by design^52–55^.

Is there a way to bridge classical, natively human-readable models that are also completely learned from data and highly performant? Classical motif analysis is limited by committing in advance to a discrete alphabet-level pattern: a consensus string, a logo, or a matrix over independent positions. We replaced this hard motif with a *soft*, or latent, motif. Every gapless window of the sequence, from a single residue up to an eight-residue stretch, is mapped by a learned linear filter into a vector that can simultaneously encode many meaningful properties. For example, identity, hydrophobicity, charge, and size can be naturally encoded in this high-dimensional latent space. A pair of two windows (each producing a “soft motif”), one from the HLA pseudosequence and one from the peptide, interact through an inner product, analogous to the way classical analysis would ask whether a particular anchor residue is compatible with a particular pocket. The model aggregates all of these pairwise compatibilities under a single learned map to produce a prediction. We call the resulting architecture *Latent Motif Interaction Aggregation* (LAMINA). The architecture is deliberately austere; it lacks nonlinearities between the motif filters and the aggregation. This constraint allows for faithful interpretability. Because the score is a bilinear form on bias-free linear features, it decomposes exactly into additive contributions from individual residue pairs, with no reference input, no surrogate, and no completeness error. The model is also small and efficient; it can be trained in roughly ten hours on a consumer desktop workstation or a few days on a consumer laptop, so the entire pipeline can be reproduced or run at low cost without a compute cluster.

In this work, we train separate LAMINA models on public binding-affinity and eluted-ligand data and benchmark them against five widely used predictors on strictly de-duplicated held-out data. We then exploit LAMINA’s interpretable architecture to compute exact residue contributions and exact, background-independent allele-specific sequence logos directly from raw logits, without Monte Carlo sampling. These model-derived explanations are compared with three complementary external references: experimentally derived affinity motifs, alanine-scanning energetics, and residue-level solvent-accessibility surface area (SASA) from experimentally determined structures.

## 2 Results

LAMINA takes two sequences: the 34-residue class I pseudosequence that encodes the polymorphic, groove-lining positions of the HLA molecule^22^, and a peptide of 8–14 residues (Fig. 1a). Each residue of each sequence is represented by a learned embedding that combines an amino-acid identity vector with a positional vector. Convolutional filters are applied over every gapless window of each sequence — every single residue, every dipeptide, and so on up to every eight-residue stretch — and produce eight latent vectors per window, one per filter. The model then exhaustively computes the inner product of every latent motif on the HLA side against every latent motif on the peptide side, producing a large matrix of pairwise motif scores (Fig. 1a, center). Each entry of this matrix represents the compatibility, as learned by the model, between one specific window of the HLA groove and one specific window of the peptide (Fig. 1d). Each pair is then weighted by a learned coefficient that depends only on which motif bank each partner came from: its unique coefficient for the two window lengths and the two filter indices (Fig. 1b). The pairwise motif map lets the model learn the significance of each soft-motif pair. Averaging the weighted compatibilities over all valid pairs yields a single scalar which a logistic transform converts to a prediction (Fig. 1c). We train one LAMINA model to predict presentation probability and another to predict a normalized binding affinity. From this design, we also show LAMINA is equivalent to a highly constrained family of length-conditioned bilinear operators (Methods).

**Figure 1:**
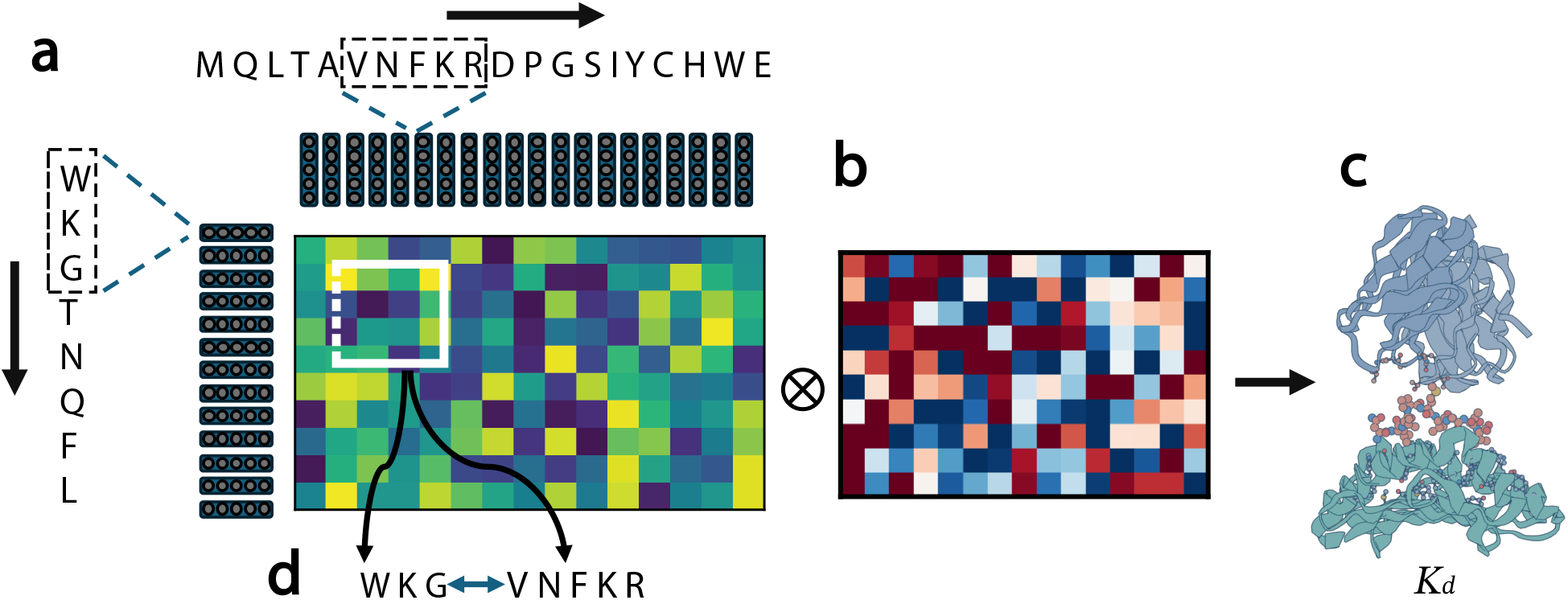
The LAMINA architecture. **a**, The HLA class I pseudosequence (horizontal) and the peptide (vertical) are embedded residue by residue. A bank of convolutional filters maps every window of length 1–8 in each sequence to a latent motif vector. Every HLA latent motif is then compared with every peptide latent motif by an inner product, giving a matrix of pairwise motif interaction scores (center; one dashed cell highlighted). **b**, A learned interaction map assigns a weight to each pair of motif banks, indexed by window length and filter identity, and is applied to the compatibility matrix element-wise. **c**, The weighted compatibilities are averaged over all valid window pairs and passed through a logistic function to give the predicted normalized binding affinity or presentation probability. **d**, Each entry of the compatibility matrix is directly readable as the learned interaction between one HLA window (here `VNFKR`) and one peptide window (here `WKG`).

### 2.1 pMHC binding-affinity prediction

We trained two LAMINA models with identical architecture and hyperparameters on the public NetMHCpan data universe, one on quantitative binding-affinity (BA) records and one on mass-spectrometry eluted-ligand (EL) records. The BA model is evaluated on a recent, prospective, independently collected set of quantitative binding measurements drawn from the Immune Epitope Database ^56,57^ (IEDB). Every peptide–pseudosequence pair appearing anywhere in the BA/EL training pool was removed from the evaluation set, which eliminated 779 of 1,382 parsed measurements and left 603 held-out measurements spanning 18 alleles. We compared against NetMHCpan-4.2, MHCflurry 2.2.1, DeepAttentionPan, ANTHEM, and TransPHLA-AOMP using a common cohort for six-model classification and for the four models with continuous affinity outputs.

LAMINA gave the highest area under the receiver operating characteristic curve at both conventional binder thresholds (Fig. 2a): AUROC 0.923 (95% CI 0.902–0.943) at 500 nM, against 0.908 for NetMHCpan-4.2 and 0.911 for MHCflurry 2.2.1. LAMINA achieved AUROC 0.908 (95% CI 0.876–0.934) at the stricter 50 nM threshold, against 0.893 and 0.889 respectively. The same ordering held in precision–recall, which is the more informative view at the low prevalence of strong binders (23.4% of records at 500 nM, 9.0% at 50 nM): LAMINA reached AUPRC 0.741 and 0.411 at the two thresholds, ahead of NetMHCpan-4.2 (0.700, 0.367) and MHCflurry 2.2.1 (0.719, 0.337).

**Figure 2:**
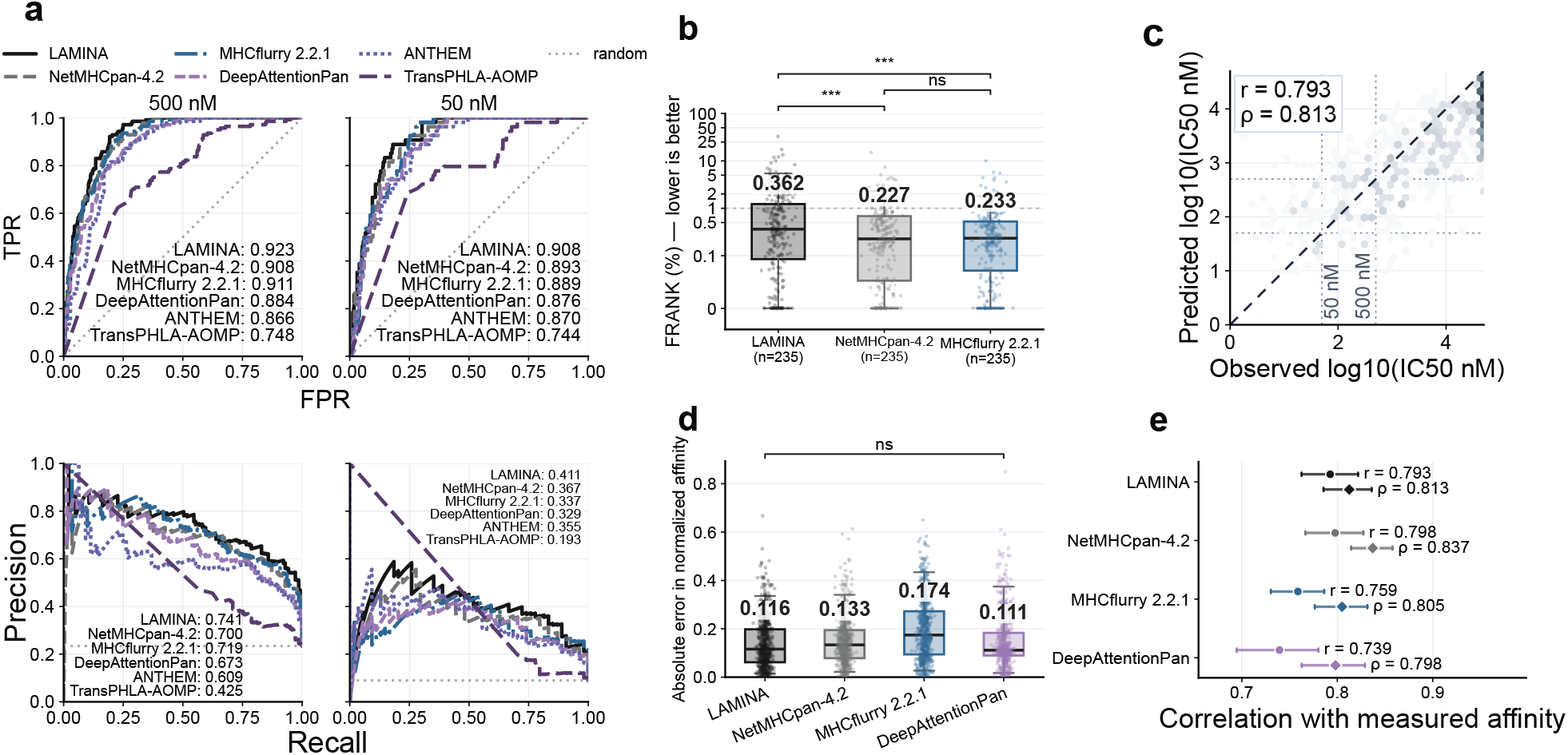
Predictive performance on held-out binding affinity and eluted-ligand recovery. **a**, Receiver operating characteristic (top) and precision-recall (bottom) curves for binder classification at 500 nM (left) and 50 nM (right) on 602 held-out IEDB measurements (141 and 54 binders respectively). Legends give AUROC (top) and AUPRC (bottom). **b**, Distribution of FRANK, the fraction of source-protein windows scored above the true eluted ligand (lower is better), over 235 Pearson *et al*. eluted ligands, shown on a symmetric-logarithmic axis, with per-epitope points, interquartile boxes, and 5th–95th percentile whiskers. Annotations give medians. Brackets show Holm-adjusted cluster-paired randomization tests over source-protein × allele clusters (^∗∗∗^*P <* 0.001; ns: not significant). **c**, LAMINA predicted affinity versus measured IC_50_ for 603 held-out measurements; the dashed line is the identity and the dotted lines mark the 50 nM and 500 nM thresholds. **d**, Absolute error in normalized affinity on the held-out cohort; annotations give medians, boxes span the interquartile range with 5th–95th percentile whiskers. **e**, Pearson *r* and Spearman *ρ* correlation with measured affinity on the same records, with 2,000-sample bootstrap 95% confidence intervals.

On the binding-affinity regression task, LAMINA’s predicted binding affinities track the measured values closely over four orders of magnitude (Fig. 2c; Pearson *r* = 0.793, Spearman *ρ* = 0.813, *n* = 603). Across models, the rank correlations are tightly clustered (Fig. 2e): NetMHCpan-4.2 attains the highest Spearman correlation (0.837, 95% CI 0.815–0.858) with LAMINA second (0.813, 95% CI 0.786–0.836), while NetMHCpan-4.2 and LAMINA have similar Pearson correlations (0.798 vs. 0.793). LAMINA achieves low calibrated error; its mean absolute error in normalized affinity units was the lowest of the models with continuous outputs (Fig. 2d; mean 0.138, median 0.116), significantly below NetMHCpan-4.2 (median 0.133, mean 0.148; paired two-sided Wilcoxon signed-rank, Holm-adjusted *P* = 3.2 × 10^−4^) and MHCflurry 2.2.1 (median 0.174, mean 0.197; *P* = 1.4 × 10^−27^), and not significantly different from DeepAttentionPan (median 0.111, mean 0.146; *P* = 0.069). In other words, LAMINA ranks binders about as well as the best existing tools and predicts how strongly they bind.

### 2.2 Recovery of eluted ligands from their source proteins

Ranking a known epitope against every peptide its source protein could have yielded is a harder and more realistic test of a presentation model than discriminating curated positives from curated negatives. We therefore used the FRANK statistic ^22,28^: for each experimentally eluted ligand, all 8–14-residue windows of its UniProt source protein^58^ are scored, and FRANK is the fraction of those windows that score above the true ligand (lower is better). We evaluated on class-I-eluted ligands from Pearson *et al*.^26^ as retrieved from IEDB. Applying the same exact peptide–pseudosequence exclusion criterion left 235 records across 208 source proteins and 20 alleles, with a mean of 4,759 competing windows per record.

LAMINA placed the true ligand in the top 0.36% of source-protein windows at the median (Fig. 2b). NetMHCpan-4.2 and MHCflurry 2.2.1 were better on this benchmark, with median FRANK of 0.227% and 0.233%. The gaps are small in absolute terms — 0.70 and 0.82 percentage points of mean FRANK — but they are consistent, and a cluster-paired randomization test that treats each source-protein × allele combination as the unit of resampling is significant (Holm-adjusted *P* = 3.0 × 10^−5^ for both comparisons), whereas the two baselines do not differ significantly from each other (*P* = 0.10). Taken together with the affinity results, LAMINA is at parity or better on binding affinity and slightly behind the specialist presentation models on source-protein ligand recovery, while remaining the only model in the comparison whose scores can be decomposed exactly.

### 2.3 Exact attributions recover binding energetics and interface geometry

By design, LAMINA’s predicted affinity logit is exactly the sum of contributions from individual HLA-peptide residue pairs. Summing that map over the HLA axis gives a single number per peptide position: the exact amount by which that residue raises or lowers the predicted binding logit, in the units of the prediction itself. Unlike generic post-hoc surrogate or Shapley-based attribution, there is no baseline input to choose, no sampling variance, and no residual to absorb ^48–50^.

As a well-established baseline, we asked what the model had learned about allele specificity using sequence logos, one of the field’s oldest readouts. For a fixed HLA allele and peptide length, LAMINA’s raw binding-affinity logit is exactly additive in the peptide’s amino-acid identities (see Methods). We therefore evaluated all 20 substitutions at each position of a 9-mer in a single batch of 180 peptide–HLA pairs and centered the resulting raw logits within each position. This produces an exact, background-independent preference logo without Monte Carlo sampling or consultation of an external ligand or motif database (Fig. 3f). Exhaustive enumeration would involve 20^9^ = 512 billion 9-mers, scaling exponentially, whereas the additive structure of LAMINA requires only 20*N* substitution evaluations for a peptide of length *N*, scaling linearly. For HLA-C*02:02, the resulting logo recapitulates the canonical motif: a dominant alanine preference at P2, a C-terminal anchor favoring tyrosine, valine, alanine, and methionine at P9, a secondary aromatic preference at P1, and comparatively weak preferences across the solvent-exposed central positions P4–P8. Across six representative alleles, the computed logos showed strong concordance with the affinity-derived IEDB SMM matrices after standardization within each allele and position (pooled Spearman *ρ* = 0.853 across 1,080 allele–position–amino-acid values) (Fig. 3e). This is a familiar sanity check of learned allele specificity rather than a novel biological finding, though it demonstrates the computational advantage of using an interpretable architecture.

**Figure 3:**
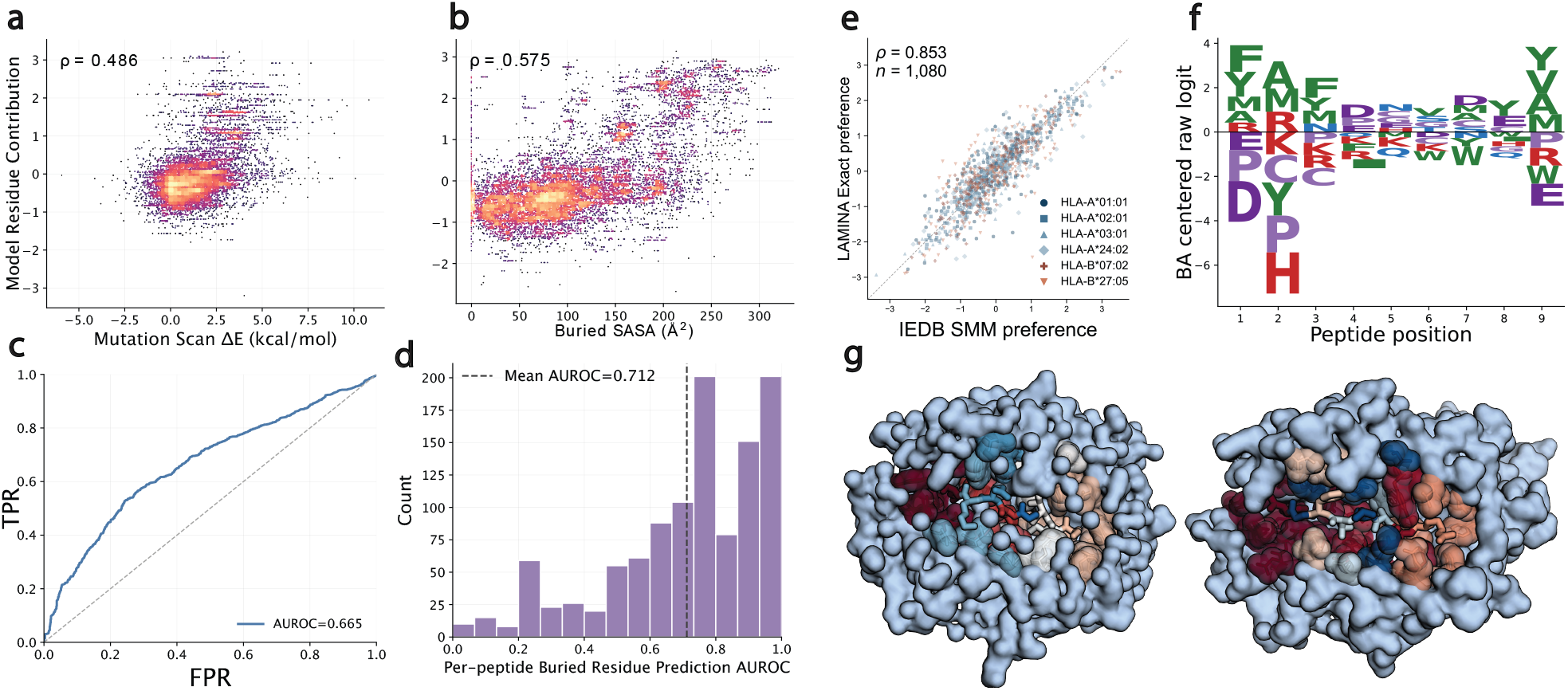
Exact attributions recover binding energetics, allele motifs and interface geometry. **a**, Exact per-residue model contribution to the predicted affinity logit versus alanine-scan interaction energy Δ*E* = *E*_mutant_ − *E*_wild type_, for 13,097 peptide single mutants across 1,413 complexes; points are colored by local density and *ρ* is the pooled Spearman correlation. **b**, Per-complex standardized model contribution versus buried solvent-accessible surface area for 14,522 peptide residues across 1,577 complexes. **c**, Pooled receiver operating characteristic for classifying poorly solvated bound residues (relative bound SASA *<* 0.5) from within-complex standardized model contribution on 14,480 residues; 42 modified residues lacking supported maximum ASA values are excluded. **d**, Distribution of the same AUROC computed separately within each of the 1,074 complexes containing both classes; the dashed line marks the mean. **e**, Agreement between exact LAMINA BA 9-mer preferences and IEDB SMM position-specific binding preferences for six HLA alleles. The 20 amino-acid scores were standardized within each allele and peptide position, yielding 1,080 allele–position–amino-acid comparisons. Colors and symbols identify alleles, the dashed line denotes equality, and *ρ* is the pooled Spearman correlation. **f**, Exact centered-raw-logit sequence logo for HLA-C*02:02, obtained from all 20 substitutions at each of nine peptide positions (180 evaluations). Letters above the axis are favored and letters below are unfavored; letter height is the magnitude of the centered raw-logit preference. **g**, Exact peptide attribution projected onto two representative pMHC complexes (PDB 3P9M, left; 7K80, right). Peptide residues are colored by their own value; each HLA pseudosequence residue takes the value of its nearest peptide residue.

We then compared interpretable attribution scores produced by LAMINA to *in silico* physical calculations. Using a curated set of 1,577 structurally characterized class I pMHC complexes from the Protein Data Bank ^59,60^, we ran alanine mutation scans in which each peptide residue was mutated to alanine (with wild-type alanine residues mutated to glycine) and the change in interface interaction energy, Δ*E* = *E*_mutant_ − *E*_wild type_, was computed after repair and rebuilding. Among available completed scans passing interface-quality filters, 13,097 single mutants across 1,413 complexes remained. Remarkably, interpretable attribution scores and mutational energy are positively correlated (Fig. 3a; Spearman *ρ* = 0.486; mean per-complex *ρ* = 0.485, median 0.517). In other words, residues to which LAMINA assigns a large positive contribution are the residues whose removal will cost the interface the most energy. Nothing in training exposed the model to a structure, a force field, or a mutation.

The same attribution scores also track interface geometry. For each coordinate-resolved peptide residue in each complex we computed the solvent-accessible surface area (SASA) buried on binding, by comparing the residue’s exposure in the HLA-bound complex with its exposure in the same peptide conformation alone (Shrake–Rupley, 1.4 Å probe)^61,62^. Across 14,522 peptide residues in 1,577 complexes, attribution and buried SASA are strongly correlated (Fig. 3b; Spearman *ρ* = 0.573; mean per-complex *ρ* = 0.593, median 0.636). Treating the problem as classification, i.e., is a residue poorly solvated while bound, defined as relative bound SASA below 50%, the within-complex standardized attribution achieves a pooled AUROC of 0.663 on 14,480 residues with supported normalization (Fig. 3c). Computed separately within each of the 1,074 complexes containing both classes, the mean AUROC is 0.710 and the median 0.750 (Fig. 3d). Projecting the one-dimensional peptide attribution onto representative structures (Fig. 3g; PDB 3P9M and 7K80) shows the expected spatial pattern, with the highest-contribution residues buried in the groove pockets and the lowest-contribution residues pointing into solvent.

## 3 Discussion

LAMINA generalizes the classical practice of mining sequence motifs and reasoning about how they meet: it learns latent motifs over every gapless window of both partners, scores every cross-partner window pair, and aggregates those scores under a single length-conditioned map. On strictly de-duplicated, held-out binding affinity, it attains the best AUROC at both the 500 nM and 50 nM thresholds and the lowest mean absolute error in normalized affinity among models with continuous outputs — not significantly different from DeepAttentionPan and significantly better than NetMHCpan-4.2 and MHCflurry 2.2.1. On source-protein recovery of eluted ligands, it is competitive but measurably behind two specialist presentation models. A model at this level of accuracy can serve as the binding-and-presentation filter in neoantigen triage, epitope prioritization for vaccine design, or repertoire-level analyses of HLA-restricted presentation. LAMINA demonstrates that for pMHC presentation and binding prediction, transparency need not be sacrificed at the cost of performance. An interpretable-by-design model does not need post-hoc explainable guarantees to be empirically demonstrated or argued; it is provably true^52^.

The most striking result is that these exact attributions correlate with real-world biochemical properties. The model was trained strictly on binding affinities and eluted-ligand labels. It never saw a coordinate file, a force field, a solvent model, or a paired mutation. Yet, the residues it assigns highest attribution to predicted binding are empirically the residues that contribute the most energy to the binding interaction, and they are also the residues that are buried on binding. The directly computed motif logos independently recover the canonical P2 and C-terminal anchors of a specific allele directly from the learned sequence model and correlate well to experimentally measured residue affinity maps. Taken together, this is direct evidence that the latent motif–motif interactions are not an arbitrary parameterization that happens to fit; the pairwise compatibilities the model learns are physically meaningful. This claim can only be made cleanly with provable exact attribution; with a post-hoc explanatory method, a correlation of this kind is confounded by the explainer. Here the quantity being correlated with Δ*E* and with buried SASA *is* the model’s prediction, split into parts that provably sum back to it, as opposed to a proxy used for post-hoc explanation.

At 4.7 million parameters, LAMINA achieves these results in about ten hours on a single desktop workstation. Interpretability did not require a larger model to compensate for lost performance; the constraints that make the model readable (linear motif extraction, positional weight sharing, exhaustive rather than learned pair selection) also make it efficient. Whole-pipeline reproduction, from data preparation through attribution, is within reach of a single researcher without cluster access.

Several limitations bound these conclusions. The held-out affinity benchmark is small because we removed every exact peptide–pseudosequence pair present in the training pool. Our exclusion criterion is defined by the shared NetMHCpan BA/EL training universe. It protects LAMINA and, largely, NetMHCpan-4.2, but the other baselines were trained on different corpora with different overlaps, and we cannot fully equalize contamination across tools that we did not train. Unknown baseline training overlaps likely favor some baselines, but their effect on relative performance cannot be quantified. Regardless, the recent reference years should in principle lower the chances of train-set overlap on the baselines. LAMINA is slightly behind the leading presentation models on FRANK.

Source-protein ranking is sensitive to the tail of the score distribution and it is possible that a presentation-specific loss, antigen-processing features, or the multi-allelic deconvolution used by NetMHCpan-4.1/4.2 and MHCflurry 2.0 would close the gap^29,30,32^. The structural analysis inherits the biases of the PDB: crystallized pMHC complexes are enriched for well-behaved, high-affinity, mostly 9-mer peptides on common alleles, and computationally derived Δ*E* values are empirical estimates with known error, which are not experimentally measured energies^63^. Finally, binding and presentation are necessary but far from sufficient for immunogenicity ^9^; nothing in this work addresses T-cell receptor recognition ^64^, and the architecture’s fine-tuning to immunogenicity endpoints is left to future work.

Beyond pMHC, the design principle is general. Many problems in molecular biology are naturally posed as “which short pattern in sequence A is compatible with which short pattern in sequence B”: transcription-factor binding sites and their cofactors, RNA-binding protein recognition elements, protease cleavage motifs, short linear motifs mediating protein–protein interactions, and antibody– antigen contacts all have this shape, and all have a tradition of hard-motif analysis that a soft-motif interaction model could generalize ^19,65^. The requirements for exactness are modest: linear feature extractors and an aggregation that is bilinear in the two sides. Thus, they are compatible with a wide range of encoders. The more interesting implication is methodological: when the interpretability is exact, the attributions become an experimental object in their own right, something that can be correlated against physics and falsified. More broadly, this work demonstrates an often true and overlooked phenomenon: often times, interpretability and performance are not tradeoffs; they can both be maximized with intentional and principled design choices.

## 4 Methods

### 4.1 Overall study design

This is a computational study with three components: (i) training a latent motif interaction model on public pMHC class I binding-affinity and eluted-ligand data; (ii) benchmarking its predictive accuracy against external predictors on held-out data from which all exact training pairs were removed; and (iii) validating the model’s exact attributions against three independent references: motif logos, alanine mutation scans, and per-residue solvent-accessible surface area computed from experimentally determined structures.

### 4.2 Model architecture

#### Inputs and embeddings

Let *A* denote the amino-acid vocabulary (22 residue symbols plus padding, class, separator, and unknown tokens; a total vocabulary size of 26). An HLA molecule is represented by its canonical class I pseudosequence *a* = (*a*_1_, …, *a*_*M*_) with *M* = 34, consisting of the groove-lining positions 7, 9, 24, 45, 59, 62, 63, 66, 67, 69, 70, 73, 74, 76, 77, 80, 81, 84, 95, 97, 99, 114, 116, 118, 143, 147, 150, 152, 156, 158, 159, 163, 167 and 171 of the mature heavy chain^22^. A peptide is *b* = (*b*_1_, …, *b*_*L*_) with 8 ≤ *L* ≤ 14. Each residue is embedded as the sum of a token embedding and a positional embedding, both learned and shared between the two sequences,

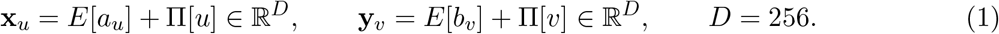

#### Latent motifs

Let ℒ = {1, 2, …, 8} be the set of motif lengths and let *R* = 8 filters be learned per length, producing *G* = |ℒ| · *R* = 64 *motif banks* per sequence. Index a bank by *g*, with length *ℓ*_*g*_ ∈ ℒ. Each bank has a filter tensor 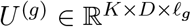 on the HLA side and 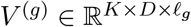 on the peptide side, with motif dimension *K* = 32. The latent motif produced by bank *g* at start position *I* is the linear map of the corresponding window,

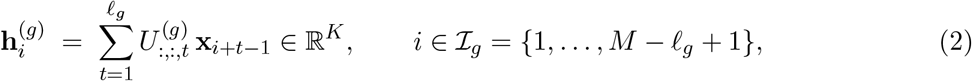

and analogously on the peptide side,

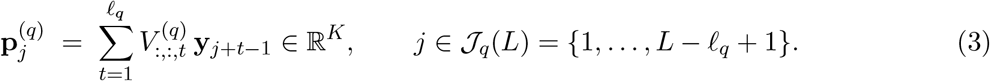

This is implemented as a multi-channel convolution. For *M* = 34 this gives ∑_*g*_ |ℐ_*g*_| = 1,952 HLA latent motifs, and for a 9-mer peptide ∑_*q*_|*J*_*q*_| = 352 peptide latent motifs.

#### Interaction and aggregation

Every HLA latent motif is compared against every peptide latent motif by an inner product. A learned motif-motif interaction map *A* ∈ ℝ^*G×G*^, whose entries depend only on the pair of motifs — i.e., only on the two window lengths and the two filter indices, allows for a learned “interaction strength” between any pair of motifs. The score is the mean of the weighted interactions:

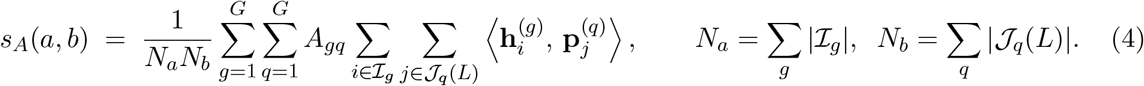

Scores are mapped between 0 and 1 with the logistic function. The affinity target is the standard normalization 1 − log(IC_50_)*/* log(50,000) with IC_50_ in nM clipped to [1, 50,000] ^20^, so normalized affinity predictions are directly comparable to published normalized affinities and can be inverted to an IC_50_ estimate.

#### Bilinear reduction

Because *A*_*gq*_ does not depend on *i* or *j*, the inner double sum in Eq. (4) factorizes. Defining length-pooled motif vectors 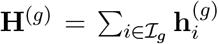 and 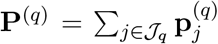, and stacking them into matrices *H, P* ∈ ℝ^*G×K*^,

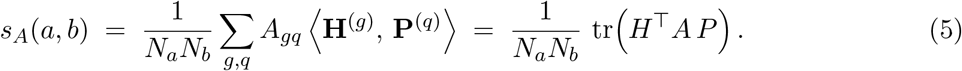

LAMINA is therefore a constrained family of length-conditioned bilinear operators on pooled latent-motif representations. This identity has two uses. Practically, it removes the need to materialize the (*N*_*a*_*N*_*b*_) interaction tensor — at the training batch size used here the dense form would require more than 10^10^ elements per micro-batch — and we exploit it in a fused Triton^66^ kernel that computes Eq. (5) directly. Conceptually, it characterizes the hypothesis class the model can express. Attributions are computed by expanding the linear filters into residue-level terms; dense and pooled scoring paths agree to numerical precision.

#### Final configuration and size

The reported models use *D* = 256, *K* = 32, *R* = 8 motifs per length, ℒ = {1, …, 8} for both sequences, *M* ≤ 34, *L* ≤ 14. The total parameter count is 4,738,048.

### 4.3 Training data

Training data were the pre-update NetMHCpan-4.2 training and evaluation snapshot distributed by DTU Health Tech, downloaded from the NetMHCpan-4.2 legacy archive. This is the broad BA/EL/IEDB/CEDAR data universe used by NetMHCpan-4.1/4.2^30^. The whitespace-delimited fold files were converted to a single JSONL table: candidate alleles for multi-allelic eluted-ligand samples were read from the supplied `allelelist` file, and every allele was mapped to its 34-residue pseudosequence through the supplied `pseudoseqs` file. Rows lacking a resolvable pseudosequence or a parseable target were dropped. The formatted table contains 17,651,353 rows: 214,804 binding-affinity (BA) and 17,349,161 eluted-ligand (EL) training rows, 42,921 IEDB and 5,172 CEDAR training rows, and 34,512 IEDB and 4,783 CEDAR evaluation rows. BA targets are the normalized affinities described above; EL targets are binary presented/not-presented labels. Multi-allelic EL rows retained all candidate pseudosequences, with one sampled uniformly on each training presentation.

### 4.4 Training methodology

Two checkpoints were trained with identical hyperparameters, differing only in which source was sampled: a BA-only model (used for all binding-affinity and interpretability analyses) and an EL-only model (used for source-protein FRANK). Restricting each model to one label type keeps the two evaluation regimes clean and avoids confounding the affinity analyses with presentation labels. Accordingly, all affinity and attribution results are produced from the BA checkpoint, and all FRANK results from the EL checkpoint.

Each run performed 15,000 optimizer steps, with an effective batch of 32,768, yielding 4.9 × 10^8^ example presentations per run. Mean-reduced micro-batch losses, including the shorter final micro-batch, were averaged during accumulation. The optimizer used was AdamW ^67,68^ with learning rate 5 × 10^−4^ decayed linearly to zero over the run, with a weight decay of 0.1, default PyTorch moment settings, and gradient-norm clipping at 1.0. Loss for the EL task was binary cross-entropy and mean squared error for the BA task. A small penalty on the mean squared latent-motif activation over valid windows (weight 10^−4^) was added to keep motif norms bounded. Data were drawn by random sample without replacement over the dataset, with each new epoch being re-shuffled. Random seeds were fixed at 0 for Python, NumPy, and PyTorch for reproducibility.

Each model was trained on an NVIDIA DGX Spark desktop workstation. Implementation used PyTorch ^69^, NumPy ^70^, SciPy ^71^ and scikit-learn ^72^.

### 4.5 Binding-affinity evaluation

#### Data

Quantitative class I binding measurements were retrieved from the IEDB PostgREST query API^56,57^ restricted to reference years 2021–2026 (1,826 rows; retrieved 2026-07-24). Rows were retained when the peptide had 8–14 residues drawn only from the 20 canonical amino acids, the allele normalized to a fully typed HLA-A/B/C four-digit specificity with a pseudosequence in the NetMHCpan table, and the quantitative measure was a positive finite exact value, with or without an “=” qualifier (inequality-qualified measurements were discarded), leaving 1,382 measurements. Retained assay annotations comprised 594 *K*_*D*_ (approximately IC_50_) measurements and nine *K*_*D*_ (approximately EC_50_) measurements; figures use IC_50_ as shorthand for the reported nM affinity. Values were converted to normalized affinity with the same transform used in training.

Any evaluation measurement whose (peptide, pseudosequence) pair occurred anywhere in the BA/EL training set was removed, eliminating 779 measurements. The final cohort is 603 measurements across 18 alleles and 80 unique peptides, of which 142 are binders at 500 nM and 55 at 50 nM. Continuous correlation and error analyses use the common four-model cohort of 603 measurements. Classification uses a common six-model cohort of 602 measurements (141 and 54 binders); one record was excluded as it uses an HLA sequence that is not supported by ANTHEM.

#### Baselines

NetMHCpan-4.2 and MHCflurry 2.2.1, with the `models_class1_presentation` weight set, were run as distributed. Their affinity outputs were used for binding-affinity classification and regression; presentation outputs were used for FRANK. DeepAttentionPan ^37^ (commit c1ed9b9e), ANTHEM ^39^ (commit 410f1153) and TransPHLA-AOMP ^38^ (commit 3ed22602) were used.

#### Metrics and statistics

Binder labels were defined at IC_50_ ≤ 500 nM and IC_50_ ≤ 50 nM. We report AUROC and AUPRC at both thresholds, Pearson and Spearman correlation between predicted and measured normalized affinity, and the distribution of absolute error in normalized affinity units. Confidence intervals for AUROC, AUPRC, and correlations are percentile intervals from 2,000 deterministic bootstrap resamples of records. Error distributions were compared with two-sided paired Wilcoxon signed-rank tests over all model pairs, with Holm correction across the full family of pairs.

### 4.6 Source-protein eluted-ligand recovery (FRANK)

#### Data

Class I eluted ligands of length 8–14 from Pearson *et al*.^26^ were retrieved from IEDB (28,408 rows), and the referenced source proteins were downloaded from UniProt ^58^. Each record was kept only when its peptide occurs verbatim in its source sequence (28,329 rows), the allele normalizes to a fully typed HLA-A/B/C specificity with a known pseudosequence, and the record is unique on (dataset, peptide, allele, accession), leaving 28,256 records. Applying the identical exact peptide–pseudosequence training-pair exclusion removed 28,021 records (99.2%). The reported cohort is 235 records over 208 source proteins and 20 alleles.

#### Protocol

For each record, candidates are all 8–14-residue windows at every position of the source protein; repeated sequences arising from repeated positions are deliberately retained, so FRANK ranks positions as in the original protocol ^22,28^. Any candidate window that forms an exact training pair with that record’s pseudosequence is removed from the candidate list, except the true peptide itself. Every candidate is scored, the true peptide’s rank is 1 +#{candidates scoring strictly higher}, and FRANK = #{strictly higher}*/*#{candidates}. The cohort averaged 4,759 candidates per record (median 3,396). LAMINA ranks candidates by raw EL logits to avoid artificial ties from sigmoid saturation.

#### Statistics

Because records from the same source protein and allele share candidate pools and are not independent, model pairs were compared with a cluster-paired model-label randomization test at the (source accession × HLA allele) level: 100,000 random sign flips of the per-cluster difference sums generate the null distribution of the mean paired FRANK difference, and the two-sided *P* value is (#{|null | ≥ | observed|}+ 1)*/*(100,000 + 1). Confidence intervals are percentile intervals from 10,000 cluster bootstrap resamples. Holm correction was applied across the three model pairs. The cohort contains 221 such clusters.

### 4.7 Exact attribution

#### Derivation

Fix an input (*a, b*) and a task map *A*. Each latent motif in Eq. (2) is a bias-free linear function of the residue embeddings, so it splits exactly into per-residue terms. Define the total contribution of HLA residue *u* to the pooled motif vector of bank *g*,

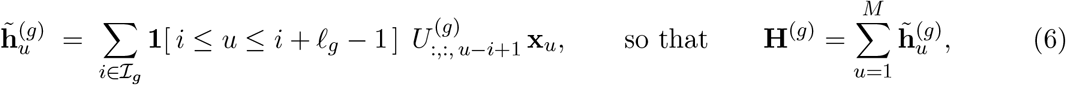

and analogously 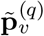 with 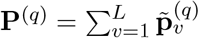. These identities simply regroup the same products.

Substituting into Eq. (5) and exchanging finite sums gives a residue-pair decomposition

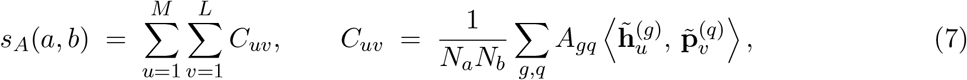

and, marginalizing over the HLA axis, a one-dimensional peptide attribution

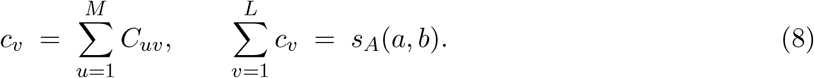

#### Relationship to gradient and post-hoc methods

Since *s*_*A*_ is bilinear in (**x, y**) and contains no bias, it is homogeneous of degree one in the peptide embeddings, and therefore 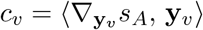 exactly. For this architecture the gradient × input attribution coincides with the peptide marginal in Eq. (8). This is exactly the property that general deep networks lack: with bias terms and elementwise nonlinearities, gradient × input has no completeness guarantee, which is why integrated gradients introduces a path integral and a reference input ^50^, why DeepLIFT introduces reference activations ^51^, why LIME fits a local surrogate whose fidelity is only empirical^48^, and why Shapley-value methods must approximate an exponential sum^49^. Each of those choices introduces a degree of freedom that can change the resulting explanation without changing the model, and their sensitivity to those choices is well documented ^73–75^. Here there is no reference, no surrogate, no sampling and no residual: the attribution is a rearrangement of the arithmetic that produced the prediction.

### 4.8 Exact centered-logit binding-affinity logos

#### Exact additive form

Fix an HLA pseudosequence *a* and peptide length *L*. Equation (7) shows that, once the HLA is fixed, the score is a linear function of the embedded peptide residues. Consequently, for each peptide position *v* there is a fixed vector

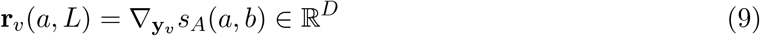

that depends on the HLA, peptide length and trained BA model, but not on the peptide’s amino-acid identities. This vector collects the contributions of every convolutional window that overlaps position *v*. The raw score is therefore

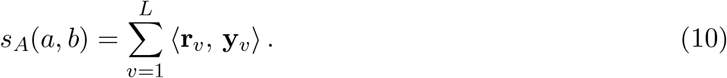

Using **y**_*v*_ = *E*[*b*_*v*_] + Π[*v*] gives

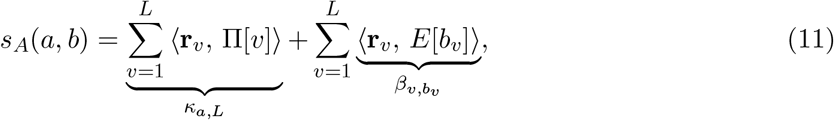

or, more compactly,

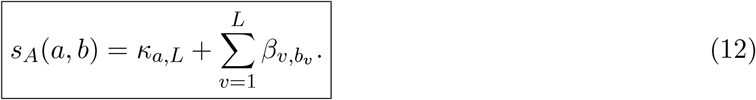

Thus overlapping motif windows remain additive after they are collected at each residue. There are no peptide–peptide terms involving products of the amino-acid identities at two different positions.

#### Exact logo values

Let *A*_20_ be the 20 canonical amino acids. For a uniform amino-acid background, the signed preference of amino acid *α* at position *v* is

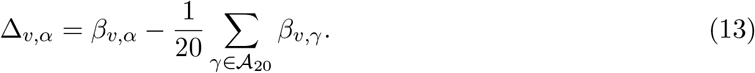

By Eq. (12), this quantity is exactly the conditional mean raw-score lift

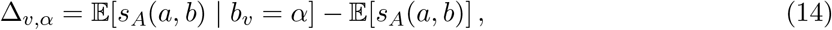

where peptide residues are sampled independently and uniformly. Contributions from every other peptide position cancel in the difference.

Starting from an arbitrary background peptide *b*^(0)^, we substituted each of the 20 amino acids at each position. Let

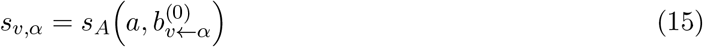

be the raw score after replacing position *v* by *α*. For fixed *v*, Eq. (12) gives

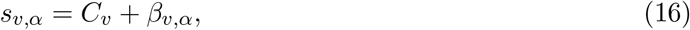

where *C*_*v*_ is identical for all 20 substitutions. Row-centering therefore removes the background exactly:

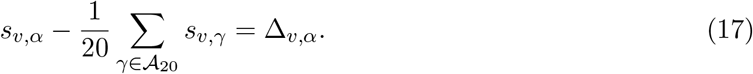

All 20*L* substitutions were evaluated in one aligned batch; a 9-mer therefore required 180 peptide– HLA evaluations. An all-alanine peptide was used as the computational background, although the centered result is independent of this choice.

Because the complete score separates by position, the globally highest-scoring peptide is also obtained exactly:

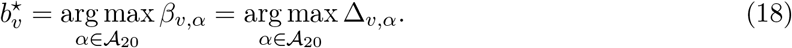

The same peptide maximizes the normalized-affinity prediction because the sigmoid is monotonic. The logo itself must nevertheless be computed from raw scores: independently centering sigmoid-transformed predictions would destroy the additive identity and make the result depend on the background peptide.

#### Visualization

HLA-C*02:02 was used for the single-allele illustration. Values were drawn as a signed, two-sided sequence logo^76^: letter height is |Δ_*v,α*_|, favored residues appear above zero, and disfavored residues appear below zero. All 20 amino-acid values were retained in the calculation; only the eight largest absolute values at each position were drawn for readability.

#### Comparison with IEDB SMM

Exact BA preference tables were additionally computed for HLA-A*01:01, HLA-A*02:01, HLA-A*03:01, HLA-A*24:02, HLA-B*07:02 and HLA-B*27:05. These were compared with the corresponding 9-mer stabilized-matrix-method matrices from the IEDB MHC-I tools archive, https://tools.iedb.org/mhci/download/. SMM coefficients contribute additively to predicted log_10_(IC_50_), for which smaller values indicate stronger binding. We therefore negated the SMM coefficients before treating them as amino-acid preferences; the sequence-independent SMM intercept was not used.

For each allele and position, the 20 exact BA values and 20 SMM values were standardized separately to zero mean and unit population standard deviation. Agreement was summarized by pooling the resulting 6 × 9 ×20 = 1,080 allele–position–amino-acid values and computing their Spearman correlation. All 20 values entered the correlation, even though only the eight largest absolute values were displayed in the logo grid. This comparison measures descriptive motif concordance and is not treated as an independent held-out performance estimate.

### 4.9 Structural cohort

The complete weekly wwPDB mmCIF archive^59,60^ (mirror of 2026-06-25) was scanned for entries containing both an MHC class I heavy-chain entity and a polypeptide entity of 8–14 residues, identified by entity description, HLA allele annotation, or heavy-chain sequence signature, yielding 1,185 candidate entries. The 1,915 previously selected chain-pair records were re-mapped by aligning each full deposited heavy-chain sequence to mature UniProt P04439 (sequence version 2; precursor residues 25–365) ^58^, using BLOSUM62, affine internal gaps (open −10, extension −0.5), and free terminal gaps. Alignments required at least 90% coverage and 70% identity across mature positions 1–182, with equally optimal alignments agreeing throughout this region and all 34 model positions present. The reference defines numbering; pseudosequences retain the deposited residues. Low-confidence and ambiguous alignments excluded 224 and 71 records, respectively, including many nonhuman MHC chains. Exact label-chain, label-sequence, author-residue and insertion-code maps were used to recompute the 34 × *L* minimum-heavy-atom distance matrices for 1,620 accepted records, of which 1,613 have a complete 34-residue coordinate mapping and 1,606 are scoreable by the model.

Two fixed structural screens were then applied. A peptide is flagged *unbound* when its closest heavy-atom distance to the entire MHC chain is at least 6 Å (25 records). A pseudosequence *pocket miss* is flagged when the peptide contacts the MHC chain within 4 Å at 20% or more of peptide positions, yet the closest pseudosequence residue is at least 5.5 Å away and pseudosequence contact is either absent or both sparse (≤ 15% of positions) and at least 2 Å more distant than the closest MHC residue (4 records). Removing both flags from the scoreable records left the analysis cohort of 1,577 complexes.

### 4.10 Alanine mutation scan

For each scanned complex, a clean two-chain PDB file containing only the HLA heavy chain and the peptide was written from the mmCIF source. FoldX^77,78^ was then run in four stages: `RepairPDB` on the two-chain complex; `AnalyseComplex` on the repaired wild type; `BuildModel` to produce one single mutant per peptide position, mutating each residue to alanine (or, for wild-type alanine, to glycine); and `AnalyseComplex` on each mutant. The quantity analyzed is Δ*E* = *E*_mutant_ − *E*_wild type_ in the FoldX interaction-energy scale (kcal mol^−1^), where positive values indicate a weakened interface.

Before analysis, interfaces were required to have a wild-type interaction energy of at most −5 kcal mol^−1^ and at least 10 wild-type interface residues, and each mutant was required to retain at least one interface residue; rows with non-finite or numerically zero energies were dropped. Of the 1,577 complexes, 99 had no completed scan and 65 were excluded by the energy filters, leaving 13,097 peptide single mutants across 1,413 complexes (925 distinct PDB entries). Each mutant was paired with the exact, unstandardized model contribution *c*_*v*_ of the mutated peptide position from the same complex. We report the pooled Spearman correlation over all mutants, together with the mean and median of the per-complex Spearman correlations.

### 4.11 Peptide solvent-accessible surface area

Residues were resolved by exact mmCIF label-chain and label-sequence identifiers, using the first coordinate model and heavy atoms, retaining modified polymer residues and consistent alternate conformers in both states. For each complex in the structural cohort, per-residue SASA was computed with Biopython’s Shrake–Rupley implementation^61,62,79^ using a 1.4 Å probe and 200 sphere points, in two states: the *bound* state, containing exactly the HLA heavy chain and its peptide, and the *free* state, containing the identical peptide coordinates alone with no relaxation. For each peptide residue,

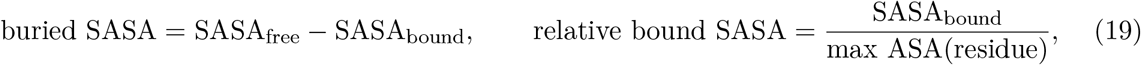

with theoretical maximum accessible areas from Tien *et al*. ^80^. Buried SASA was clipped at zero and relative bound SASA to [0, 1]. This is a purely geometric calculation and involves no energy model. The 90 peptide positions without resolved coordinates were excluded from SASA analysis. The analysis set is 14,522 residues over 1,577 complexes. The 42 modified residues lacking supported maximum ASA values were retained for buried-SASA analysis but excluded from normalized-SASA analyses (a maximum ASA value is required to compute normalized-SASA), leaving 14,480 residues.

Because the absolute scale of the attribution varies between complexes, contributions were standardized within each complex (zero mean, unit standard deviation across that peptide’s positions) for the SASA analyses; the FoldX analysis above uses unstandardized values. We report the pooled Spearman correlation between standardized contribution and buried SASA, the mean and median of the per-complex correlations, and a binary analysis in which a residue is positive when its relative bound SASA is below 0.5. For the binary target we report the pooled AUROC over residues with supported normalization and the distribution of AUROCs computed separately within each complex that contains both classes (1,074 complexes).

For the structural projections in Fig. 3g, each peptide residue is colored by its own value and each HLA pseudosequence residue is colored by the value of its nearest peptide residue.

## Data availability

All data used in this study are public. The NetMHCpan-4.2 training and evaluation folds are available from DTU Health Tech. Binding-affinity, eluted-ligand and epitope records are available from the IEDB (https://www.iedb.org) and CEDAR; the eluted-ligand benchmark derives from Pearson *et al*.^26^. Source protein sequences are from UniProt ^58^ and structures from the wwPDB^60^. FoldX is available from the FoldX Suite and is not redistributed here.

## Code availability

The complete implementation is available at our GitHub repository which can be found at https://github.com/nyuolab/LAMINA.

## Author contributions

S.F.C. developed and implemented the computational models, performed model training and evaluation, and conducted computational experiments and analyses. R.J.S. and E.K.O. contributed to the interpretation of the modeling results, the preparation, validation and documentation of the code used in this study, and to the writing and revision of the manuscript.

